# Automatic processing of numerosity in human neocortex evidenced by occipital and parietal neuromagnetic responses

**DOI:** 10.1101/2020.07.15.203786

**Authors:** Amandine Van Rinsveld, Vincent Wens, Mathieu Guillaume, Anthony Beuel, Wim Gevers, Xavier De Tiège, Alain Content

## Abstract

Humans and other animal species are endowed with the ability to *sense*, represent, and mentally manipulate the number of items in a set without needing to count them. One central hypothesis is that this ability relies on an automated functional system dedicated to *numerosity*, the perception of the discrete numerical magnitude of a set of items. This system has classically been associated with intraparietal regions, however accumulating evidence in favor of an early *visual number sense* calls into question the functional role of parietal regions in numerosity processing. Targeting specifically numerosity among other visual features in the earliest stages of processing requires high temporal and spatial resolution. We used frequency-tagged magnetoencephalography (MEG) to investigate the early automatic processing of numerical magnitudes and measured the steady-state brain responses specifically evoked by numerical and other visual changes in the visual scene. The neuromagnetic responses showed implicit discrimination of numerosity, total occupied area, and convex hull. The source reconstruction corresponding to the implicit discrimination responses showed common and separate sources along the ventral and dorsal visual pathways. Occipital sources attested the perceptual salience of numerosity similarly to both other implicitly discriminable visual features. Crucially, we found parietal responses uniquely associated with numerosity discrimination, showing automatic processing of numerosity in the parietal cortex, even when not relevant to the task. Taken together, these results provide further insights into the functional roles of parietal and occipital regions in numerosity encoding along the visual hierarchy.

**Significance Statement:** Approximating the number of items in a set has been identified as a building block of mathematical cognition but the processing of numerosity is not fully understood. The natural correlation between numerosity and other visual features makes it difficult to test whether the number of items is a perceptual primitive or whether it needs to be recombined at a higher level. We used frequency-tagged magnetoencephalography to localize the implicit discrimination of numerosity within the visual hierarchy. We found that numerosity yielded occipital responses, supporting that the human visual system can grasp it at a single glance. Crucially, numerosity also yielded specific parietal responses, showing that numerosity is a perceptual primitive with a unique automatic involvement of parietal cortex.

## Introduction

Posterior parietal cortex has been recurrently associated with the processing of numbers and numerical magnitudes, especially regions along the intraparietal sulcus (IPS). Numerical magnitude can be grasped under different formats (symbolic, i.e., 10, verbal, i.e., *ten*, or non-symbolic, i.e., ·········). Dehaene and colleagues (2003) advanced the idea of a core representation of numerical magnitude in an abstract, modality-independent format located in the IPS. Neuroimaging evidence corroborated this idea (e.g. Piazza et al., 2004; see Sokolowski et al., 2017 for a meta-analysis) and further highlighted a spatial gradient of non-symbolic number representations along the dorsal stream, quite similarly to the low-to high-level visual processing gradient along the ventral visual stream (Roggeman et al., 2011). Within this occipito-parietal stream, an increasing decoding of numerosity was identified, which was peaking at parietal regions, supporting the functional role of these regions for the highest level representation of numerosity (Eger et al., 2009; Bulthé et al., 2014). Further, ultra high-field functional magnetic resonance imaging (fMRI) allowed mapping numerosity coding in the parietal cortex with exquisite spatial details (Harvey et al. 2013, 2015).

In parallel, a set of adaptation studies by Burr and Ross (2008, 2017) led to the *visual number sense* hypothesis, which rather focused on the early visual processing in numerosity extraction. Accordingly, numerosity could be encoded in primary visual cortices because it exhibits adaptation properties similar to other primary visual features (e.g., color). Neuroimaging evidence supported occipital cortex involvement in numerosity extraction, corroborating the possibility of an early decoding of numerosity information in the visual hierarchy. The observation of parietal regions in numerosity processing was suggested to be linked to the behavioral magnitude judgment task (e.g., Lasne et al. 2011). Some passive viewing experiments recorded only occipital but no parietal activity associated to unattended numerosity decoding and thus postulated that the IPS could rather support attentional mechanisms linked to numerosity processing (DeWind et al., 2018; Castaldi et al., 2019).

The role of the primary visual cortex for numerosity has also been emphasized by electrophysiological studies disclosing early neural sensitivity to numerosity (Park et al., 2015, 2017). More specifically, frequency-tagging electroencephalography (EEG) allowed recording steady-state visual responses evoked specifically by both numerosity and some continuous visual features demonstrating implicit discrimination of these features (Guillaume et al, 2018; Van Rinsveld et al. 2020). This approach distinguishes responses to different features based on their presentation frequency, without the need to isolate them in the visual presentation, which is crucial because numerosity is intrinsically correlated with non-numerical, continuous magnitude parameters (Norcia et al., 2015). However, the low spatial resolution of EEG did not allow clearly disentangling occipital and parietal activity, which is essential to further characterize the occipital/parietal functional dissociation in the processing of numerosity and non-numerical magnitudes along the dorsal pathway.

The current study used MEG to combine the exquisite time resolution necessary for high-frequency tagging with sufficient spatial resolution to enable source localization along the visual hierarchy. We tested the hypothesis that implicit discrimination of numerosity and continuous magnitudes occurs in the visual occipital cortices. We measured to what extent parietal sources contribute to implicit discrimination to test the attentional account of IPS role in numerosity extraction. This was achieved using the fast periodic visual stimulation paradigm illustrated in Figure 1. We presented arrays of dots that randomly varied in numerosity and continuous features. Similarly to oddball paradigms, one feature was held constant across standards but was varied deterministically at the rate of 1.25 Hz (deviant items, occurring every eight items). The feature identified by deviant items was either numerosity (i.e., the number of dots) or a continuous magnitude (i.e., dot size, total area, dot density, and convex hull). Importantly, both standard and deviant stimuli within each block differed because only one feature was periodically fixed (e.g., numerosity) while all others randomly varied. We expected that frequency-tagging neuromagnetic responses would allow specifically discriminating the periodic features and locating the underlying neocortical sources.

**Figure 1.**
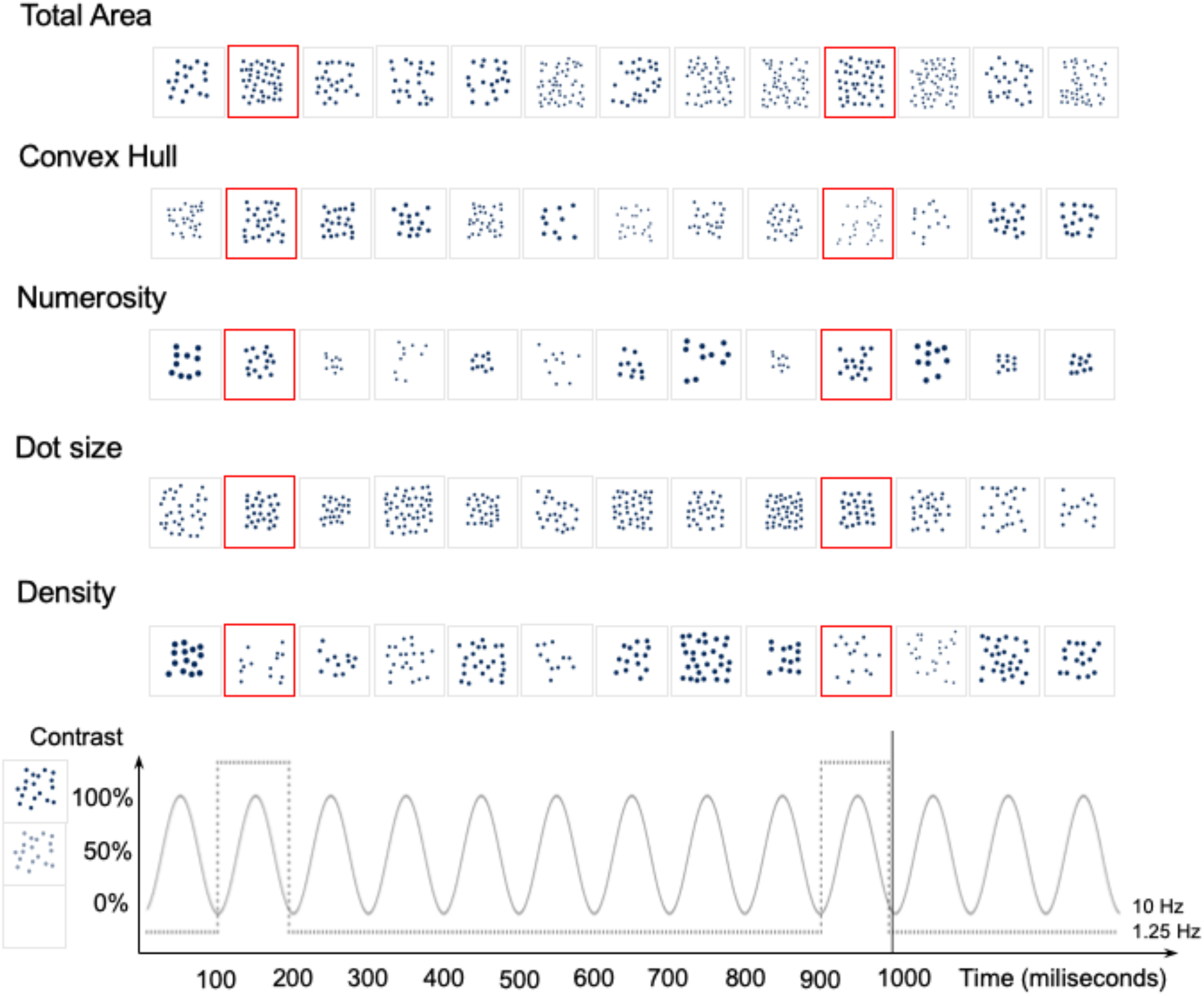
Sequences of dot patterns varying five features: dot size, total area, convex hull, density, and numerosity. Upper panels depict an example of 1.3 s-long time series (13 stimuli) for the five experimental conditions. In each condition, the relevant feature is held constant across standard stimuli (grey borders) and changed only in the deviant stimuli (red borders) presented periodically, all other features being varied randomly. Ten stimuli are presented each second (10 Hz presentation rate), and deviants occur every eight item (1.25 Hz oddball frequency). Each visual stimulus was presented with gradual contrast modulated sinusoidally at 10 Hz, as shown in the lower panel.

## Methods

### Participants

Twenty-one healthy adults participated in this study. Volunteers suffering from, or with a history of, neurological or neuropsychological disorders, learning disability such as dyscalculia, or uncorrected visual impairment, were not included. We excluded one participant due to unusable structural MRI necessary for MEG source localization. The final sample thus consisted of twenty participants, with a mean age of 23.5 years (range 20–29 years, 12 females). We followed APA ethical standards to conduct the study, which received prior approval by the CUB – Hôpital Erasme Ethics Committee (reference: P2018/362). The entire experiment lasted 3 to 4 hours in total, and participants received 10 € per hour for their participation.

### Visual stimulation and apparatus

Pictures showing dot arrays were generated with NASCO dot generation toolbox (Guillaume et al., 2020) as in previously validated EEG work (Van Rinsveld et al., 2020) (for examples, see Figure 1). We generated 100 standard pictures and 100 deviant pictures for each condition, each 850-by-850 pixel. Deviants were designed as standards except that the magnitude of one feature (total area, convex hull, numerosity, dot size, density) was increased by a factor of 150%. Previous evidence showed that this ratio is easily discriminated when adults perform magnitude judgment tasks (Barth, 2003). From this database of pictures, we generated sequences of 440 stimuli made of series of 7 standards followed by one deviant. To avoid periodicity in the stochastic fluctuations in the non-relevant features, we extracted the Fourier spectrum of the time series for each feature presentation and z-score normalized it. We only retained sequences in which the spectral z score of the feature of interest exceeded 2.32 (corresponding to the unilateral threshold at 99% of a standard normal probability distribution) at 1.25 Hz deviant frequency, and concurrently the spectral z score of the other features was below this threshold. The sequence of each condition was repeated four times to increase the signal-to-noise ratio.

The fast periodic presentation of the stimuli was handled with the Psychophysics Toolbox (MATLAB, The MathWorks, Brainard, 2017; Kleiner et al., 2007). Visual stimulation was displayed at 1 meter from the participants at the centre of a MEG-compatible screen inside the magnetically shielded room (Maxshield™, MEGIN, Helsinki, Finland), via a DLP projector (Panasonic PT-D7700E, New Jersey, USA; 60 Hz refresh rate, 1366 × 768 pixels of resolution) placed outside the room and projection through a feedthrough.

### Experimental procedure

Participants were comfortably seated in the MEG armchair in front of the screen. They were instructed to look at the screen and to keep their gaze fixed on a diamond that was continuously displayed in blue at the centre of the screen. Each time the fixation diamond changed colour (red), participants had to press a key on a MEG-compatible response pad. Changes randomly occurred between four and eight times in a sequence. This aimed at maintaining a similar vigilance level across conditions and refraining participants to look away. Stimulus sequences were displayed with sinusoidal contrast modulation from 0% to 100% (Lochy et al., 2017) (Figure 1) at the base rate of 10 Hz (one stimulus presented every 100 ms), for a total duration of 44 s length. Five sequences were presented, in which one feature (total area, convex hull, numerosity, density, and dot size) was held fixed among standards and systematically changed in deviants, which occurred every eight items (1.25 Hz oddball frequency). Two seconds of fade-in and two seconds of fade-out were added respectively at the beginning and at the end of each sequence to ensure smooth transition to the stimulations but were discarded from analyses. The order of the sequences was randomly counterbalanced across participants. At the end of the experiment, no participant reported noticing neither periodicity nor the feature of interest.

### Data acquisition

Visual evoked magnetic fields were measured using a whole-scalp-covering MEG system (Triux™, MEGIN, Helsinki, Finland) containing 102 triplets of sensors, one magnetometer and two orthogonal planar gradiometers. Neuromagnetic activity was recorded continuously during each sequence, with internal active compensation (Maxshield™, MEGIN, Helsinki, Finland), analog band-pass filtering between 0.1 and 330 Hz, and digital sampling at 1 kHz. The timing of sequence start and end (i.e., without fade-in and fade-out periods) was identified by a trigger signal. Four coils on the participants’ head allowed to track head position continuously. Their location with respect to fiducials and over 300 scalp points sampling the head shape were obtained by three-dimensional digitalization (Fastrak Polhemus, Colchester, VT, USA). High-resolution structural 3D T1-weighted MRI of the participant’s brain was acquired after the MEG session with a 1.5 T MRI scanner (Intera, Philips, The Netherlands).

### Data processing

Environmental noise and head movements were corrected off-line using signal space separation (Taulu et al., 2005) as implemented in the Maxfilter™ software (MEGIN, Helsinki, Finland, v2.2). Independent component analysis was then applied on the resulting MEG signals band-filtered between 0.5 and 45 Hz, to identify and suppress ocular and cardiac artefacts (Vigaro et al., 2000). The cleaned data were chunked into four 44 s-long epochs corresponding to the repetition of the same visual sequence and were averaged to increase signal-to-noise ratio (Liu-Shuang et al., 2014).

To enable the reconstruction of source activity underlying MEG data, we also processed the structural MRIs to compute individual forward models. MRI and MEG data were first coregistered manually using the digitized fiducial points for initial approximation and head-surface points for refinement (MRIlab, MEGIN, Helsinki, Finland). Forward modeling was then performed using the single-layer boundary element method implemented in the MNE-C suite (Gramfort et al., 2014), based on MRI segmentation obtained from the Freesurfer software (Fischl, 2012). The forward model was computed for three-dimensional sources located on the nodes of a volumetric brain grid, which was built from a regular 5-mm grid in the Montreal Neurological Institute (MNI) template MRI (16102 nodes) and transformed into individual MRIs using a non-linear spatial normalization (Ashburner & Friston) implemented in the SPM12 toolbox (Friston et al., 2007).

### Spectral analysis

The Fourier coefficients of the averaged MEG data chunk were computed via discrete Fourier transformation of their first 40 seconds, leading to a frequency resolution of 0.025 Hz, and source projected by minimum norm estimation (MNE) (Dale et al., 1993). The MNE projector was built from the individual forward model, noise covariance obtained from empty room MEG recordings, and a regularization parameter set via the prior consistency condition (Wens et al., 2015). Spectral amplitude was finally obtained at each source location as the Euclidean norm of the three components Fourier magnitudes.

Given our oddball paradigm, we focused on the detection of spectral peaks at the base (10 Hz) and oddball (1.25 Hz) frequencies and their harmonics. We extracted the amplitude spectra in frequency intervals centered on the frequency of interest and some harmonics (base frequency: 10 and 20 Hz; oddball: all multiples of 1.25 Hz smaller than 10 Hz) and summed them (sum-based amplitude, SBA). To assess the size of the peak at the target frequency, we compared the SBA for the target frequency bin to the SBA for neighboring bins (10 adjacent bins on both sides, the closest left and right neighbors being discarded). This comparison was carried out by standardizing the target SBA value, i.e., by subtracting the mean SBA over neighbor bins and dividing by their standard deviation. This led to two brain maps of standardized SBA per participant, one corresponding to the base rate and the other to the oddball rate.

Statistical significance of these maps was assessed at the group level using unilateral parametric *t* tests against the null hypothesis that there was no difference. The significant threshold at *p* = .05 was *t*_19_ = 1.725. However as each map encompassed a large number (i.e., 16102) of tests, a substantial number of false positives would be expected. The family-wise error rate was controlled by estimating the number of spatial degrees of freedom in MNE maps based on the forward model rank (*N*=62 in this data, see Wens et al., 2015 for details) and applying Bonferroni correction, i.e., performing all univariate tests at *p* < 0.05/*N* = 0.0008). The resulting corrected threshold was *t*_19_=3.632.

To compare the conditions that showed significant oddball responses, we ran contrast analyses by subtracting conditions two by two and generating statistical maps assessing the significance of the difference at the group level using bilateral parametric *t* tests against the null hypothesis that all *ts* = 0. The significant threshold at *p* = .05 was *t*_19_ = 2.086, and the family-wise error corrected threshold was *t*_19_=3.930. These contrast maps are reported to visualize the discrepancies between conditions at the whole brain level but not to accurately localize the sources of these differences. Indeed, recent evidence demonstrated that MEG parametric contrast maps are suited to assess the existence of differences between conditions but not to draw conclusions about the source localization of these differences (Bourguignon et al., 2018). Analyses on non-contrast maps should be preferred to identify the sources of the observed differences. To overcome this limitation, we further ran repeated-measure analyses of variance to assess the differences of SBA between conditions in the sources that were the maximum of each condition as reported in Table 1. ANOVAs were computed with JASP software (JASP, 2018). Bonferroni-Holm corrections for multiple comparisons were applied for post hoc comparisons between conditions.

**Table 1.** Sources localization of the significant oddball responses for the five conditions. X, Y, Z, *t*: MNI coordinates of the local maxima in the *t* maps, and the corresponding *t*_19_ value. Voxels: number of suprathreshold voxels in the connected cluster of the corresponding local maximum. Labels: provided for comparison with topographical maps shown in Figure 3.

| **Condition** | **Peak Localization** | **X** | **Y** | **Z** | **$t$** | **Voxels** | **Labels** |
| --- | --- | --- | --- | --- | --- | --- | --- |
| Total Area | Right primary visual cortex | 30 | -82 | 10 | 5.20 | 211 | A |
|  | Left intraparietal sulcus | -30 | -72 | 60 | 4.45 | 5 | B |
| Convex Hull | Right lateral occipital | 45 | -77 | 15 | 7.39 | 514 | A |
|  | Right superior temporal gyrus | 55 | -62 | 20 | 4.23 | 28 | B |
|  | Left Fusiform gyrus | -25 | -42 | -15 | 4.05 | 9 | C |
| Numerosity | Right supplementary motor area | 15 | -17 | 50 | 4.46 | 9 | A |
|  | Right intraparietal sulcus | 30 | -47 | 45 | 4.04 | 10 | B |
|  | Left precuneus | -10 | -72 | 60 | 3.99 | 2 | C |
|  | Right cuneus | 20 | -87 | 45 | 3.97 | 4 | D |
|  | Right middle temporal gyrus | 65 | -52 | 10 | 3.93 | 2 | E |
| Density | Right superior parietal lobule | 15 | -62 | 75 | 3.67 | 1 | / |
| Dot size | No source above significance threshold |  |  |  |  |  | / |

## Results

At the base rate of 10 Hz, SBA peaks were observed in all conditions, with maximal peak location in the medial occipital regions (map maximum *t*=10.73, averaged over the five conditions) and were located in the occipital regions (Figure 2). These steady-state responses at the base rate did not attest any discrimination of the feature of interest, as expected from our experimental design.

**Figure 2.**
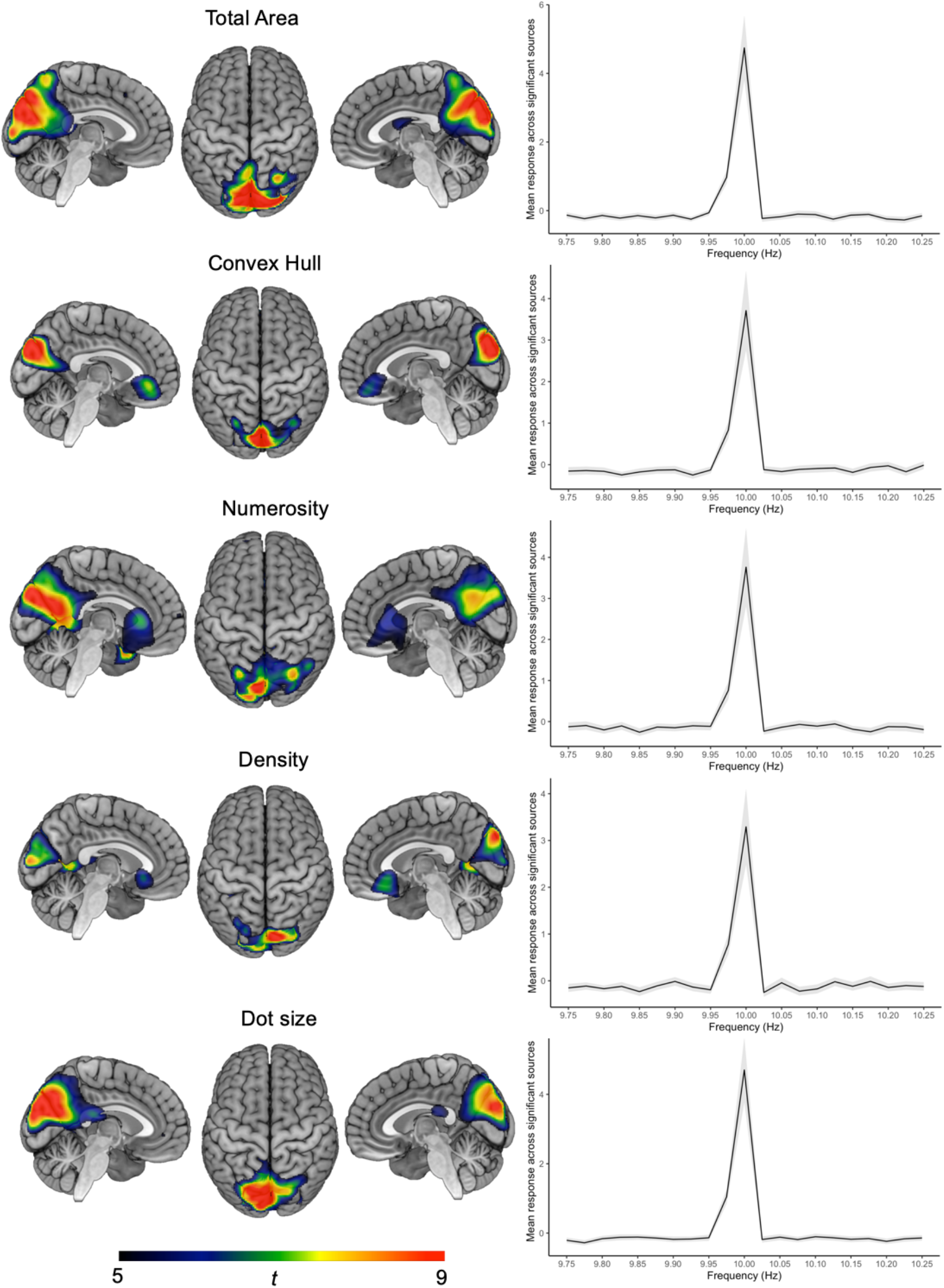
**Statistical maps of standardized SBA at the base rate (i.e., 10 Hz)** for the five conditions (total area, convex hull, numerosity, dot size and density). The color scale corresponds to the statistical *t*_19_ values. Graphs represent the mean standardized SBA response across significant sources as a function of frequency. Grey ribbon depicts standard deviations across these sources.

At the oddball rate (1.25 Hz), significant SBA peaks emerged clearly when modulating the magnitude of three features only: total area, convex hull, and numerosity, with different source locations (see Figure 3 and Table 1).

**Figure 3.**
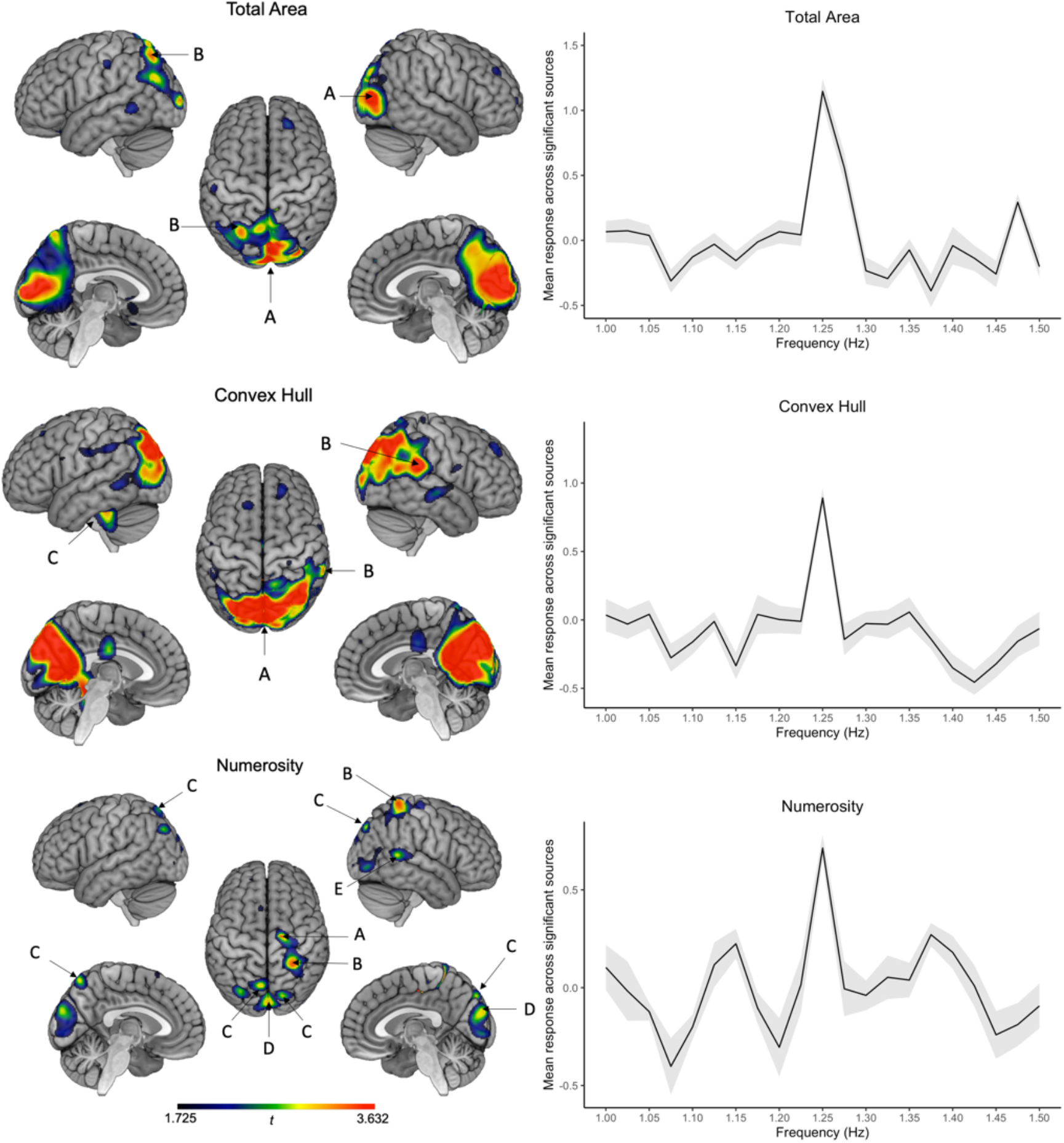
**Statistical maps of standardized SBA at** the seven first harmonics of 1.25 Hz for the three conditions where significant periodic oddball responses were recorded (total area, convex hull, and numerosity). Color scale corresponds to *t*_19_ values: *t*(uncorrected)= 1.725, *t*(corrected)=3.632. Labels correspond to distinct sources, as summarized in Table 1. Graphs represent the mean standardized SBA response across significant sources as a function of frequency (the seven first harmonics of each frequency on the x axis were considered in the mean responses). Grey ribbon depicts standard deviations across these sources.

The sources of the oddball response to periodic changes in total area were located in bilateral medial occipital regions with a local maximum in the right primary visual cortex. A second source was located at the left intraparietal sulcus. Changes in convex hull identified bilateral medial occipital regions, with a local maximum at the right lateral occipital gyrus. A second source emerged at the right superior temporal gyrus, and a third more ventrally, along the left fusiform gyrus.

Changes in numerosity (i.e., the number of dots) disclosed right occipital and right temporal sources, as in the previous condition, and a source in the left precuneus. Two other close but distinct sources emerged, one in the right supplementary motor area and the other in the right intraparietal sulcus.

Density periodic modulations only identified a single-voxel supra-threshold source in the right superior parietal lobule, which is not visible on source maps because of smoothing. Finally, no significant peak was observed at the oddball frequency in response to the dot size condition.

To compare the level of synchronization on the oddball frequency between conditions, we ran repeated-measure ANOVAs with condition (3) as within-subject factor in each maximum peak identified from the previous analysis (see Table 1). Statistical maps of the contrasts are presented in Figure 4 for visualization purposes and the results from the ANOVAs are summarized in Figure 5.

**Figure 4.**
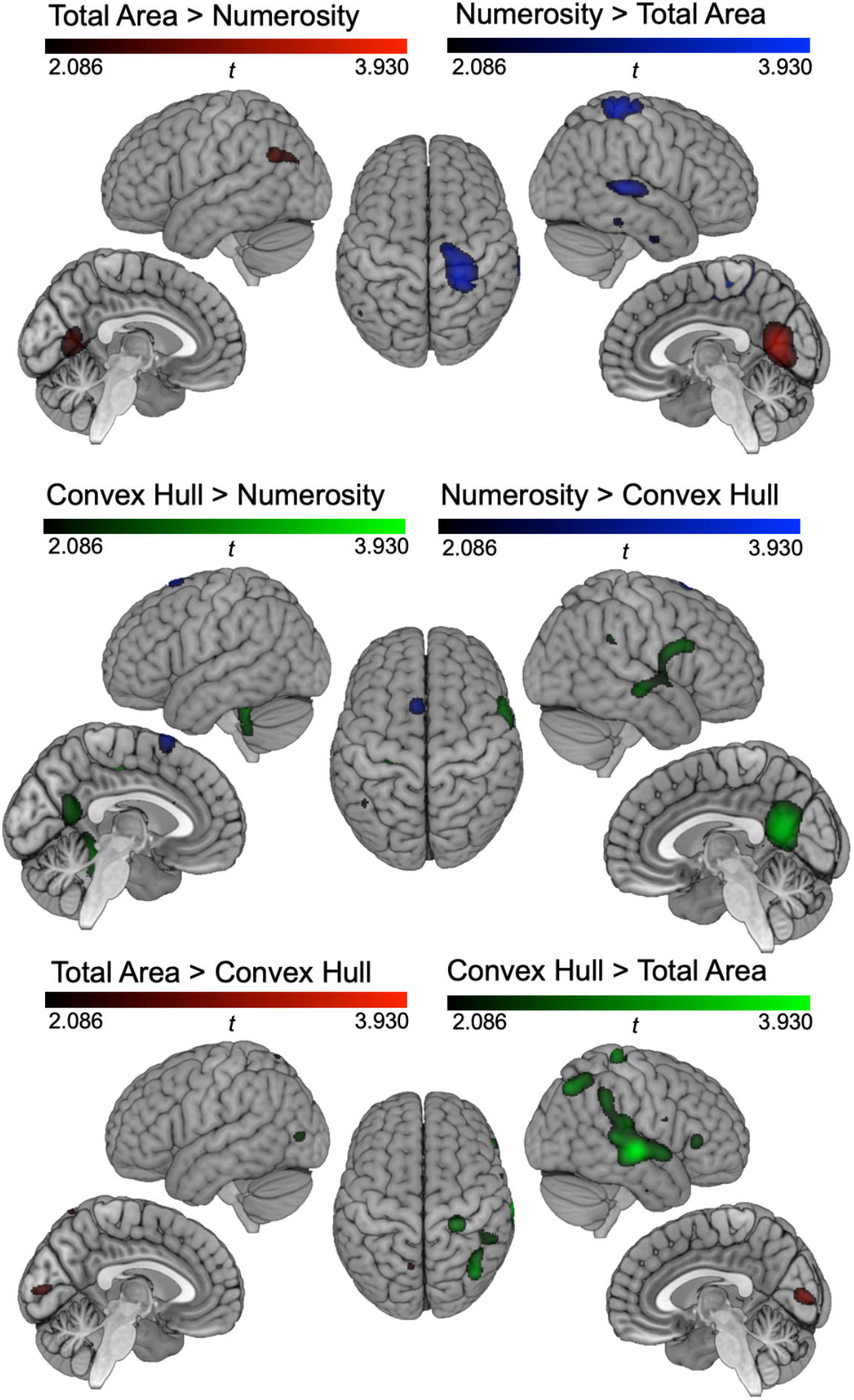
Statistical maps of the contrasts between the three conditions where significant periodic oddball responses were recorded (total area, convex hull, and numerosity). Color scale corresponds to *t*_19_ values: *t*(uncorrected)= 2.086, *t*(corrected)=3.930.

**Figure 5.**
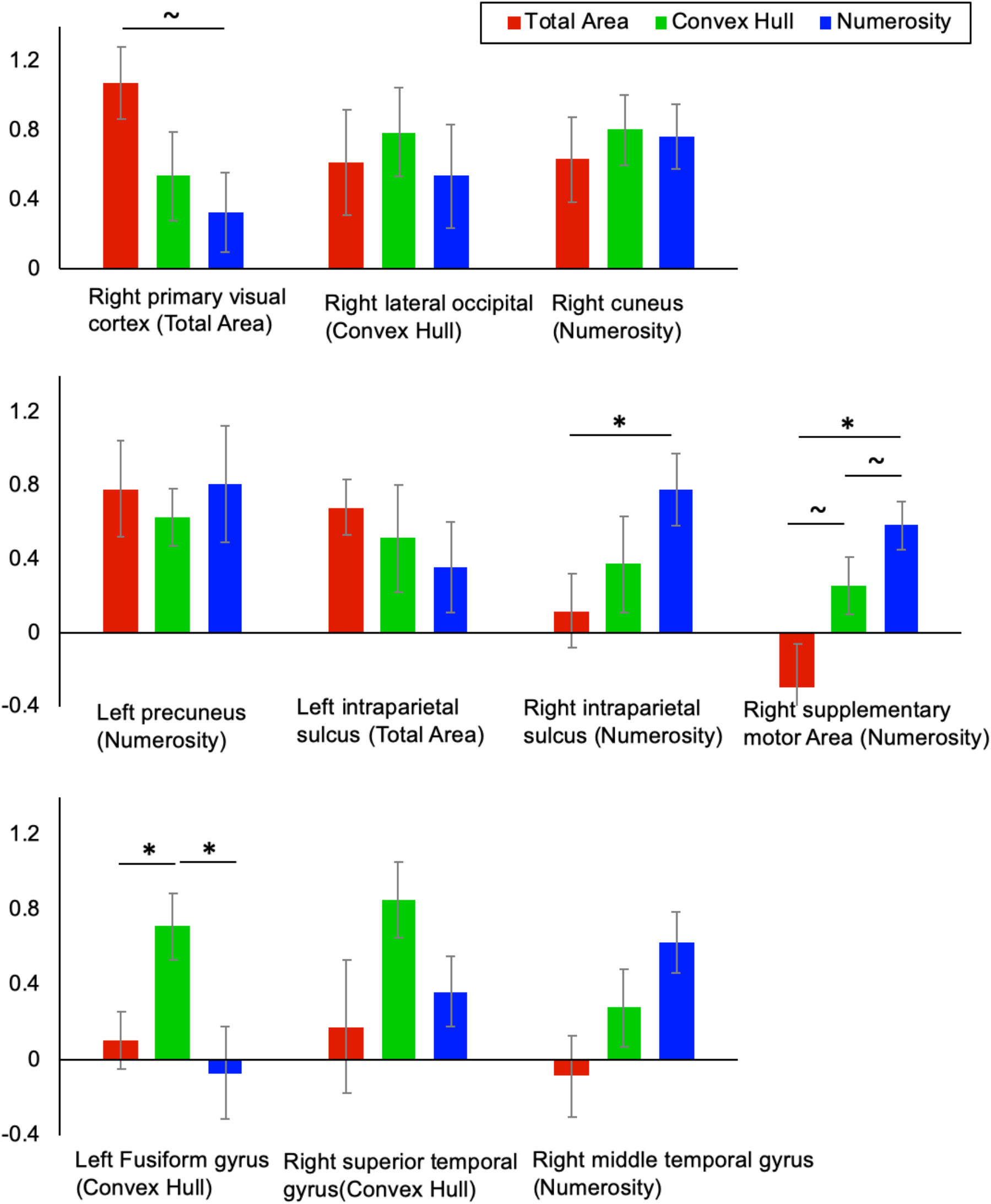
Mean SBA as a function of the local maximum peaks from each condition. The condition in which each peak was identified as a maximum is mentioned under brackets. Color categories depict the SBA at the oddball rate for the Total Area (red), Convex Hull (green), and Numerosity (red) conditions. The upper, middle, and lower graphs gather together sources located respectively in occipital, in parietal and frontal, and in temporal regions. Significant differences between conditions are marked by a star. Error bars represent standard errors from the means.

At the right primary visual cortex peak from the Total Area condition, we observed a marginal effect of condition, *F* (2,38) = 2.537, *p* = .092, η^2^ = .118. Post-hoc comparisons showed that SBA of Total Area was larger than that of Numerosity, *t*(19) = 2.157, *p* = .022, while other post-hoc condition comparisons did not reach significance, *t*(19) < 1, *p*s > .2. For both right occipital peaks for Convex Hull and Numerosity, no effect of condition was observed, *F* (2,38) = 0.268, *p* > .2, η^2^ = .014 and *F* (2,38) = 0.221, *p* > .2, η^2^ = .012.

Concerning central and parietal peaks, both the left precuneus maximum from Numerosity and the left intraparietal sulcus maximum from Total Area showed no effect of condition, *F* (2,38) = 0.521, *p* > .2, η^2^ = .027 and *F* (2,38) = 0.177, *p* > .2, η^2^ = .009. At the right supplementary motor area maximum from Numerosity, there was an effect of condition on the SBA, *F* (2,38) = 6.118, *p* = .005, η^2^ = .244. Post hoc comparisons revealed a larger SBA in Numerosity than in Total Area, *t*(19) = −3.219, *p* = .014, a marginal difference between Convex Hull and Total Area, *t*(19) = −2.158, *p* = .087, but no difference between Convex Hull and Numerosity, *t*(19) = −1.412, *p* = .174. Moreover, at the right intraparietal sulcus peak from Numerosity, there was a marginal effect of condition, *F* (2,38) = 2.921, *p* = .066, η^2^ = .133. Post hoc comparisons showed that Total Area and Numerosity differed significantly, *t*(19) = −3.109, *p* = .017, while other comparisons did not reach significance, *t*(19) < 1, *p* > .2.

In temporal regions, an effect of condition was observed at the left fusiform gyrus peak from Convex Hull, *F* (2,38) = 4.950, *p* = .012, η^2^ = .207. Post hoc comparisons showed a difference between Total Area and Convex Hull, *t*(19) = −3.015, *p* = .021 and between Convex hull and Numerosity, *t*(19) = 2.761, *p* = .025, but not between Total Area and Numerosity, *t*(19) < 1, *p* > .2. At the right superior temporal gyrus maximum from Convex Hull, there was no significant effect of conditions, *F* (2,38) = 1.691, *p* = .198, η^2^ = .082. At the right middle temporal gyrus maximum from Numerosity, the condition had a marginal effect, *F* (2,38) = 2.744, *p* = .077, η^2^ = .126, though post hoc comparisons did not show any significant differences (Total Area vs. Numerosity, *t*(19) = −2.230, *p* = .114, Convex Hull vs. Numerosity, *t*(19) = −1.622, *p* > .2, and Total Area vs. Convex Hull, *t*(19) = −1.009, *p* > .2).

In summary, the ANOVAs comparing the three conditions yielding significant oddball synchronization corroborate the differences pictured in the contrast maps. The occipital sources located in the lateral occipital and cuneus regions are relevant to the three conditions with a tendency toward a more stronger response of the primary visual cortex for the Total Area condition. Some of the parietal sources seem equivalent across conditions: left precuneus and intraparietal sulcus, while the right intraparietal sulcus and the right supplementary motor area are found only for Numerosity and Convex Hull with a tendency to a stronger response in Numerosity than in Convex Hull. Concerning the temporal regions, the fusiform gyrus response seems to be a specific of the Convex Hull condition as it is not found in the other conditions. Further right temporal gyrus sources were found both in Convex Hull and Numerosity but not in Total Area, although statistical comparisons between conditions in these regions were not significant.

## Discussion

Steady-state visual evoked neuromagnetic responses were used to identify the neural correlates of the encoding of numerosity and of continuous magnitudes. Results showed significant frequency-tagged neural responses to the deviant Numerosity, Total Area and Convex Hull. Source reconstruction highlighted the respective involvement of common and distinct regions in implicit discrimination of numerical and continuous magnitude. Primary visual cortex was the most prevalent source for total area, while further parietal and temporal regions were the more crucial in numerosity and convex hull encoding respectively. Importantly, the right IPS was specifically relevant to numerosity extraction even in an implicit discrimination requiring no explicit task and even when numerosity cannot be derived from other visual parameters. These results thus discard the pure top-down attentional function of the IPS in numerosity processing.

These results thus show the robustness of implicit discrimination of numerosity, total areal and convex hull in MEG (Van Rinsveld et al., 2020). Crucially, the source reconstruction shows that steady-state responses to periodic changes of numerosity are generated by early visual regions, but also parietal regions and to a lesser extent by temporal regions. Similarly, steadystate responses to periodic changes of total area and convex hull were generated by a combination of similar regions, although at respective contribution of these sources differed from those contributing to numerosity discrimination. This supports the hypothesis that decoding of numerosity and of certain continuous magnitudes occurs very early in the dorsal visual pathway as some sources of the recorded responses were located in occipital cortex for both numerical and non-numerical magnitudes. Our results thus support the visual number sense hypothesis (Burr & Ross, 2008), i.e., the idea that numerosity can be processed as a primary visual feature similarly to color or luminance.

The neural mechanism behind numerosity encoding has been described as following by the accumulator model: a first layer of neurons handles the sensory input detection, a second layer proceeds to input normalization and summation, and a last layer implementing number selection (Verguts et al. 2004). The current data support an early summation process that would occur both in parietal and occipital cortex and interestingly not only for numerosity but also for continuous magnitude. Crucially, the implicit discriminations observed here could not be performed based on the location of the dots or other parameters as the design ensured random variation of other parameters across the sequences.

In the literature both early visual cortex and parietal regions supporting numerosity processing have been reported, depending on the experimental paradigm and on the type of measures. Particularly, activation of parietal regions was reported to be modulated by attention to the numerosity (Castaldi et al., 2019). An fMRI metanalysis contrasted brain activations from studies using active discrimination tasks and passive viewing of non-symbolic stimuli (Sokolowski et al., 2017): Passive designs still comprised brain activations covering the right precuneus, superior parietal lobule and middle occipital gyrus. A visuo-spatial account of these findings was proposed because the superior parietal lobule is specifically associated with visuo-attentional processing involved in non-symbolic numerical tasks. The current study provides further evidence of parietal activity in absence of any explicit magnitude discrimination task and even in the absence of any conscious perception of the periodic changes (i.e., no participant reported noticing periodicity or dimension changes in any experimental condition).

Moreover, we also observed a temporal source for the Convex Hull condition along the fusiform gyrus that was not present in both other conditions. This region is usually associated to the ventral stream of object recognition and in this case could contribute to the automatic discrimination of the changes in the object shape (Martin et al., 2007; Liu et al., 2008). Indeed, considering the collection of dots holistically as a whole object in itself, its convex hull delimits the shape occupied by this object in the visual scene (Watson et al., 2014). Taken together, we thus provide evidence in favor of an automatic processing of numerosity and continuous magnitude, even without paying attention to the dimensions, discarding the pure top-down attentional account for parietal regions.

The observed source discrepancies between the three conditions yielding implicit discrimination responses have important implications for the theoretical model of an *Approximate Number System*, i.e. a distinct functional system for numerosity encoding that is specific for numerosity and does not generalize to all kind of magnitude extraction (Dehaene et al., 2003; Walsh, 2003). Previous evidence in favor of ANS were not reflecting a pure distinction between numerosity and continuous magnitude extraction because of the strong correlations between both and were complicated by the fact that processing both types of magnitude might partially share a common neural basis (Bueti & Walsh, 2009). Although there are some commonalities between the three conditions showing that some steps may be similar, also major discrepancies emerged in the sources generating the implicit discrimination responses. The current results thus attest the involvement of a pattern of brain responses that is specific to involuntary numerosity processing and that is functionally distinct from a general system that would process all kind of magnitude similarly.

Some authors argued in favor of a summation coding of numerosity in the parietal regions that would be related to the spatial disposition of the dots in the visual scene (Cavdaroglu & Knops, 2019). Their claim is based on evidence that decoding of numerosity from activation patterns in those regions is only observed for simultaneous presentation of the dot arrays in a comparison task, by opposition to sequential presentations of dot arrays where numerosity could be only decoded from occipital regions along the calcarine sulcus. In contrast, the current study encompassed both parietal and occipital sources related to the discrimination of numerosity, though we only used sequential presentations. Our results thus support the existence of a dedicated coding for numerosity both in parietal and occipital regions even in case of sequential presentation of the stimuli. This corroborates evidence of a functional dissociation between magnitude and spatial coding of numerosity among the intraparietal regions (Kanayet et al., 2018), showing that neither numerosity nor continuous magnitude coding can be reduced to sole spatial coding.

Posterior parietal regions comprise several nodes of the dorsal attention network. Specifically, intraparietal regions are typically associated to spatial attention (Silver & Kastner, 2009). These regions receive visual input from the primary visual areas but also through the superior colliculus route, and they influence the activity of primary visual regions in return. Attention orientation can be segregated in two distinct mechanisms: exogenous orienting of attention that is the involuntary orientation toward a salient stimulus due to the stimulus itself (bottom-up), and endogenous stimulus-driven attention where attention is reoriented to a stimulus that is relevant to a particular task (top-down) (Chica et al., 2013; Bisley et al. 2011). The neuromagnetic responses to the fast periodic stimulation used in the current study are likely to be driven by the former involuntary attentional mechanisms but do not preclude that in magnitude judgment tasks that have been used extensively across the literature, a mixture of both types of orientation of attention co-exists and interact. Indeed, IPS, IPL and SPL activity are modulated by the endogenous visuospatial attention and leading to reorientation of attention adapted to the demand of the task (Kastner et al. 1999; Corbetta et al. 2000). Previous evidence showed that posterior parietal cortex is involved both in conscious and non-conscious processing of visual stimuli (Kravitz et al., 2011).

The implicit discrimination of magnitudes observed through the frequency-tagged responses generated by both occipital and parietal regions suggests that the system processes automatically some features of the visual scene linked to the number of elements and other global magnitude features (Greene & Oliva, 2009). We demonstrated here that these isolated features are salient even in a task-irrelevant context. Crucially, the system also involuntary keeps track of those features across time ensuring that the next stimuli processing will take into account some characteristics of its predecessors. These results could thus be framed in the larger scope of the predictive coding theory which states that the mind is organized hierarchically to minimize prediction error with a constant feedback from higher-level regions that adjusts the predictions of lower-level regions in order to make optimal inferences about the environment (Friston, 2008; Ester et al., 2016). The current study suggests that the dorsal visual stream can handle an efficient general description of the scene combining very early decoding of numerical and continuous magnitude information with a dynamic adjustment of perceptual experience.

In conclusion, the early visual regions would be able to discriminate numerosity and some of the continuous magnitudes (total area and convex hull) and the parietal regions may support the persistence of the information over short timescales. The frequency-tagged neuromagnetic responses provide evidence in favor of an automatic feature-based attention spontaneously directed towards numerosity and some continuous magnitude properties related to the whole visual scene. Crucially the experimental design ensured that the observed discrimination responses were invariant both to spatial disposition of the dots and to the intrinsic correlations among these dimensions because of the strict control of the visual stimulation.

## Author contributions

AV, AC, and WG designed the research, AV and AB performed the research, AV and MG contributed unpublished reagents/analytic tools, AV, VW, and XD analyzed the data, AV, VW, XD, and AC wrote the paper.

## Acknowledgement

This research was funded by the European Union’s Horizon 2020 research and innovation program under the Marie Skłodowska-Curie grant agreement No 799171 to AV and by PDR project No T. 1052.15 of Fonds National de la Recherche Scientifique to AC and WG. The authors declare no conflict of interest that might be interpreted as influencing the research, and APA ethical standards were followed in the conduct of this work.

